# Metabolomics of Thermoregulatory Transitions in the Eastern Skunk Cabbage (Symplocarpus foetidus)

**DOI:** 10.64898/2026.05.23.727417

**Authors:** Johanna Toth, Riley Mantulak, Maya Bozzo-Rey, Alexander G. Little

## Abstract

The Eastern skunk cabbage (*Symplocarpus foetidus*) is a thermogenic plant with remarkably precise thermoregulatory control. Its unique sexual dimorphism in thermogenic strategy, homeothermy in female-phase flowers versus diurnal heterothermy in male-phase flowers, offers a powerful natural framework to identify thermoregulatory signaling pathways. We used untargeted metabolomics and LC-MS/MS to compare metabolite profiles across four groups: homeothermic females sampled at midday and midnight, and diurnally heterothermic males sampled at midday and midnight. Pairwise comparisons and cross-filtering against circadian- and sex-specific metabolite changes yielded 190 candidate thermoregulatory metabolites. Principal component analysis revealed that thermogenic state (warm vs. cold) was the primary axis of metabolic variation, accounting for nearly 55% of variance. Metabolite set enrichment analysis identified significant enrichment for octadecanoid formation from linoleic acid, linoleic acid oxylipin metabolism, and eicosanoid metabolism via cyclooxygenases. Oxylipins including 9-oxoODE, 9(S)-HOT, 13(S)-HOT, and 15-OxoETE were among the compounds most strongly associated with low thermogenic activity. We propose that lipoxygenase-derived cyclopentenone oxylipins may suppress thermogenesis by inhibiting thioredoxin o (TRXo), thereby inactivating the alternative oxidase (AOX) pathway that drives heat production. These findings suggest convergence between plant and mammalian thermoregulatory pathways, with linoleic acid-derived oxylipins and prostaglandins emerging as conserved regulators of metabolic heat production across deeply divergent lineages.

## Introduction

The evolution of endothermy lies at the heart of some of the most exciting and contentious debates in biology (see Koteja, 2004; Kemp, 2006; Lovegrove, 2017; Lovegrove and Seymour, 2019). Endothermy is most emblematic of birds and mammals, but it also occurs in a diverse array of taxa across the animal kingdom (e.g., Heinrich, 1972; Standora, 1982; Heinrich, 1987; Block et al., 1993; Wegner et al., 2015), and even throughout eukaryotes more broadly (e.g., Meeuse, 1975; Watling et al., 2008). This diversity has served as a cornerstone for exploring broader themes in physiology, specifically by providing an evolutionary framework to explore the mechanisms that underlie convergence in complex physiological adaptations (Block and Finerty, 1993; Little et al., 2010; Grigg et al., 2022).

The capacity to maintain a stable body temperature in dynamic thermal environments (i.e., homeothermy) requires that metabolic heat production (i.e., thermogenesis) is coupled to dynamic feedback integration, like a biological thermostat. At a coarse level, there is phylogenetic convergence in the mechanisms that drive thermogenesis itself (e.g., upregulation of aerobic metabolism via mitochondrial uncoupling; Pörtner, 2004; Wagner, 2008; Block, 2011). However, the regulatory pathways that control ‘biological thermostats’ are not well resolved beyond hypothalamic integration in more traditional avian and mammalian models. Understanding the signaling pathways that regulate homeothermy across diverse organisms can offer valuable insights into the evolutionary origins and consequences of endothermy (e.g., Little and Seebacher, 2014; Hirose et al., 2019).

Thermogenic species of plants are relatively rare but occur across several families (e.g., Araceae, Cycadaceae, Nymphaeaceae, Annonaceae, and Nelumbonaceae). Eastern skunk cabbage (*Symplocarpus foetidus*; Araceae) is particularly unique in that it blooms in the early spring in Eastern North America, even as snow and ice cover the ground and temperatures still routinely plummet below freezing (Knutson, 1974). Like other thermogenic flowers, the Eastern skunk cabbage is protogynous (i.e., maturation of female reproductive organs first; Seymour & Schultze-Motel, 1997). Its flowers are bisexual, where they transition from the female phase for fertilization to the male phase for pollen release (Seymour & Schultze-Motel, 1997; Ito-Inaba et al., 2009A). The Eastern skunk cabbage is exceptional in its thermoregulatory capacity: it remains homeothermic for days to weeks, rather than just a few hours (Knutson, 1974). Throughout the female stage (approximately 1-2 weeks), spadix temperatures are sustained at approximately 23°C, even as ambient temperatures drop as low as -14°C (Seymour and Blaylock 1999; Seymour, 2001; Ito-Inaba et al., 2009B). Notably, as the plant transitions to pollen production (i.e., through the bisexual then male phase), it adopts a diurnal heterothermic strategy, where the spadix sustains approximately 23°C during the day but drops to only a few degrees Celsius above ambient temperatures overnight (Ito-Inaba et al., 2009B). This shift in thermogenic activity, which gradually diminishes from the top of the spadix to the bottom, suggests that thermogenic strategies may vary according to reproductive needs (Ito-Inaba et al., 2009A).

As in animals, plant thermogenesis depends on futile cycling to enhance rates of mitochondrial respiration for metabolic heat production. Respiration rates in the spadices of the Eastern skunk cabbage are remarkably high, even exceeding those of many endothermic animals (Knutson, 1974; Seymour and Blaylock, 1999; Holdredge, 2000). Here, thermogenesis is facilitated by the expression of both alternative oxidase (AOX) and an uncoupling protein (UCP) in the inner mitochondrial membrane (Seymour et al., 2010; Seymour, 2001; Onda et al., 2007). AOX is an alternative terminal oxidase found in the electron transport systems (ETS) of many plant and animal taxa (Juszczuk and Rychter, 2003; McDonald et al., 2009) that facilitates a bypass of the classic cytochrome oxidase pathway (Seymour, 2001). Unlike cytochrome oxidase, however, AOX does not contribute to the proton motive force (Onda et al., 2007), effectively uncoupling oxidative phosphorylation to ramp up mitochondrial respiration (Onda et al., 2007; Seymour, 2001). The broad role of AOX in thermogenic plants can be considered analogous to uncoupling protein 1 (UCP1) in mammals, where heat production is favoured over ATP synthesis. The thermoregulatory precision of the Eastern skunk cabbage is also reminiscent of endothermic vertebrates, suggesting that intermediate metabolic regulators may feedback to fine-tune rate-limiting pathways (Seymour, 2004).

Mediators of thermogenesis have not yet been identified in the Eastern skunk cabbage. Salicylic acid was originally proposed as the primary thermoregulatory signalling molecule in thermogenic plants (Raskin et al., 1987; Raskin et al., 1990), though this proposed role is somewhat contentious (Seymour, 2004). Despite experimental effects of SA on AOX expression and respiration, the gradual fluctuations of SA concentrations in Eastern skunk cabbage spadices appear incompatible with the plant’s rapid thermoregulatory responses when ambient temperatures change, characterized by lag times of only 20-30 min (Seymour, 2004). Thyroid hormones play critical thermoregulatory roles in birds and mammals (see Colin et al., 2005). While plants are generally not renowned for their repertoire of thyroid hormones, iodine is highly reactive and elevated levels in the environment can facilitate thyroid hormone synthesis simply as a by-product of metabolism (e.g., phytoplankton and seaweed; Chino et al., 1994; Heyland, 2004). Intriguingly, iodine content of the Eastern skunk cabbage has long been noted among the highest of terrestrial plants (Marine et al., 1933), which potentially suggests that thyroid hormone or its analogues (i.e., iodothyronines or iodinated tyrosines) may occur in the Eastern skunk cabbage.

In addition to classic signalling molecules, Onda & Ito (2005) also observed decreases in sucrose, glucose, and fructose levels in post-thermogenic Eastern Skunk Cabbage, indicating that reduced heat production is linked to lower substrate levels. Similarly, Koyamatsu et al. (2023) found that the Japanese Skunk Cabbage (*Symplocarpus renifolius*) upregulates sugar transporters (STPs) and pyruvate production to stimulate AOX activity during thermogenesis. In particular, pyruvate promotes AOX activity when supplied in high concentrations (Millenaar et al., 2003). Thus, thermoregulatory control may also be exerted simply through the regulation of fuel availability.

In this study, we took a metabolomics approach to investigate thermoregulatory signalling pathways in the Eastern skunk cabbage. We leveraged the unique sexual dimorphism in endothermic strategy (i.e., homeothermy in female flowers vs. heterothermy in male flowers) to identify potential thermoregulatory metabolites. Specifically, we conducted comparative metabolomics between *(i) heterothermic male flowers at night (cold)* and *homeothermic female flowers at night (warm)*, and *(ii) heterothermic male flowers at night (cold)* and *homeothermic male flowers at midday (warm)*. This design allowed us to identify metabolites correlated with thermoregulatory activity, where orthologous comparisons with *homeothermic female flowers at midday (warm)* provided an intrinsic control for the confounding effects of sex and circadian rhythm across sample groups. Determining the circuitry that underlies thermoregulatory phase shifts in the Eastern skunk cabbage is important to understand the regulation behind what the, arguably, most basic biological thermostats. More broadly, this work explores whether intracellular thermoregulatory signaling pathways may be conserved across the tree of life. Identifying such pathways may uncover novel (or analogous but overlooked) thermoregulatory molecules in animals, particularly humans. Plant polyphenols, for instance, already represent a rapidly growing class of compounds to target thermogenesis for obesity and other metabolic syndromes (Boccellino and D’Angelo, 2020; Zhang et al., 2021; Wang et al., 2022).

## Methods

### Temperature monitoring and tissue sampling

We worked on a population of Eastern skunk cabbage (> 500 individuals) in Southwestern Ontario, Canada (Grey Sauble conservation wetland). Spathes of 20 plants were carefully opened with a scalpel and the sexual stage of the spadix was classified as female (N=12), bisexual (N=3), or male (N=5) according to Barriault et al (2021). To characterize thermal patterns in this population, spadix temperatures were recorded using an advanced thermal imaging camera (FLIR E54) in 6 hour-increments over 24 hours, beginning at 11 am in April 7-8, 2022 (Fig. 1). Male and bisexual spadices were collapsed into a ‘non-female’ category, owing to their similar heterothermic states.

**Figure 1.**
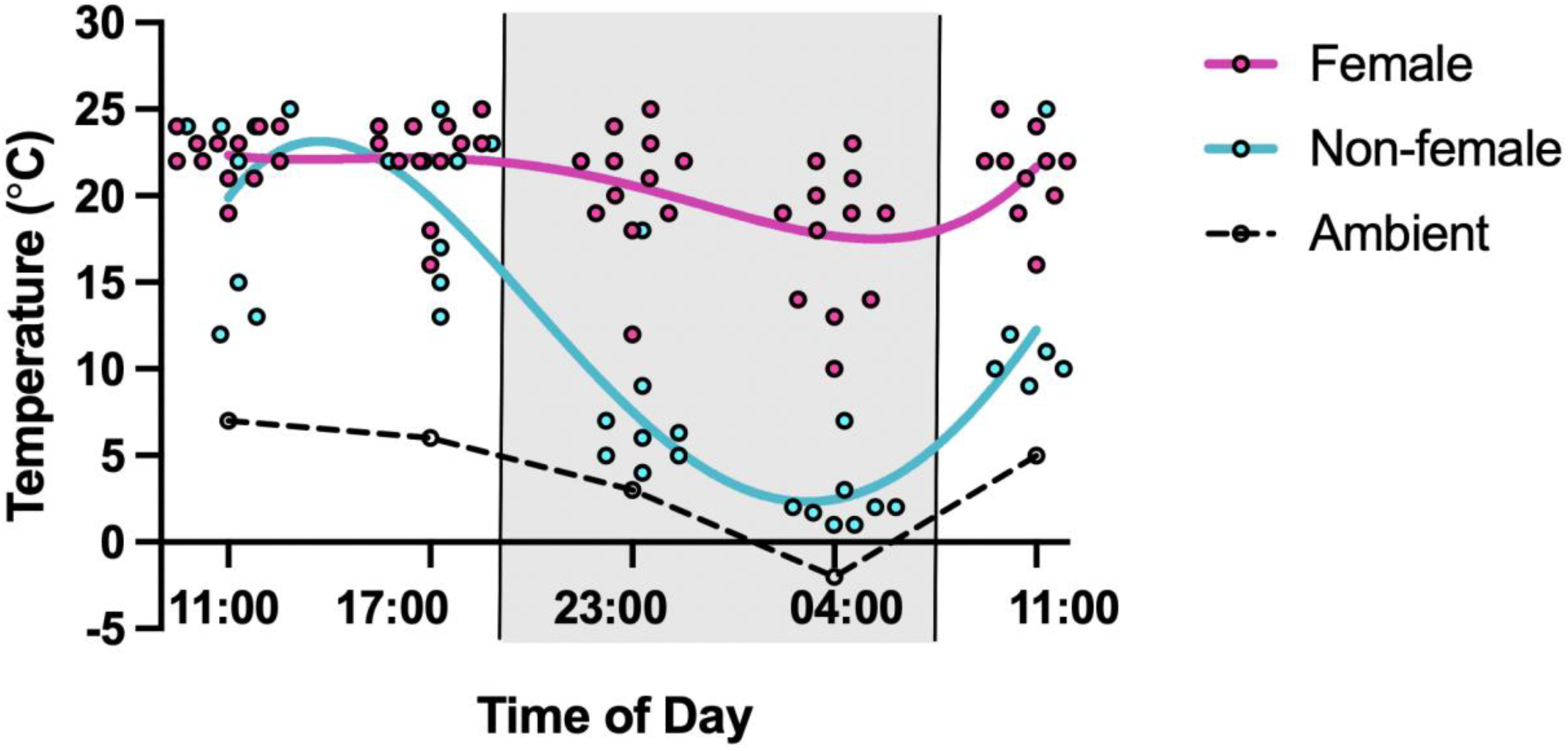
Thermal patterns in female (red) and non-female (blue; male, bisexual) spadices over a 24-hour period. Females (N=12) maintain 20-25°C spadix temperatures across the 24 hours, whereas non-female flowers (N=8) maintain 20-25°C spadix temperatures during the day but drop to only approximately 2-5°C above ambient temperature (black dashed line) during the night (grey shading).

We returned to the same population of Eastern skunk cabbage to collect spadix tissues sampled either at midday (April 7) or midnight (April 8), 2023. For plants sampled at midday (11:00-13:00), we carefully sectioned the spathes with a scalpel to reveal the spadix. We then recorded the sexual phase of the spadix (Fig. 2a,b) and its temperature using an advanced thermal imaging camera (FLIR E54). Spadices were amputated and sectioned longitudinally before being flash frozen in liquid nitrogen and stored at -80°C until later analysis. For plants sampled at midnight (23:00-01:00), we first identified, labelled, and recorded spadix temperatures at midday. We then returned to these individuals at midnight for thermal imaging and spadix collection as described above. This experimental approach (Fig. 2c) allowed us to specifically target plants at the onset of the male reproductive phase, where they are homeothermic during the day and heterothermic at night, as opposed to those nearing the end of the male reproductive phase, where thermogenesis ceases altogether.

**Figure 2.**
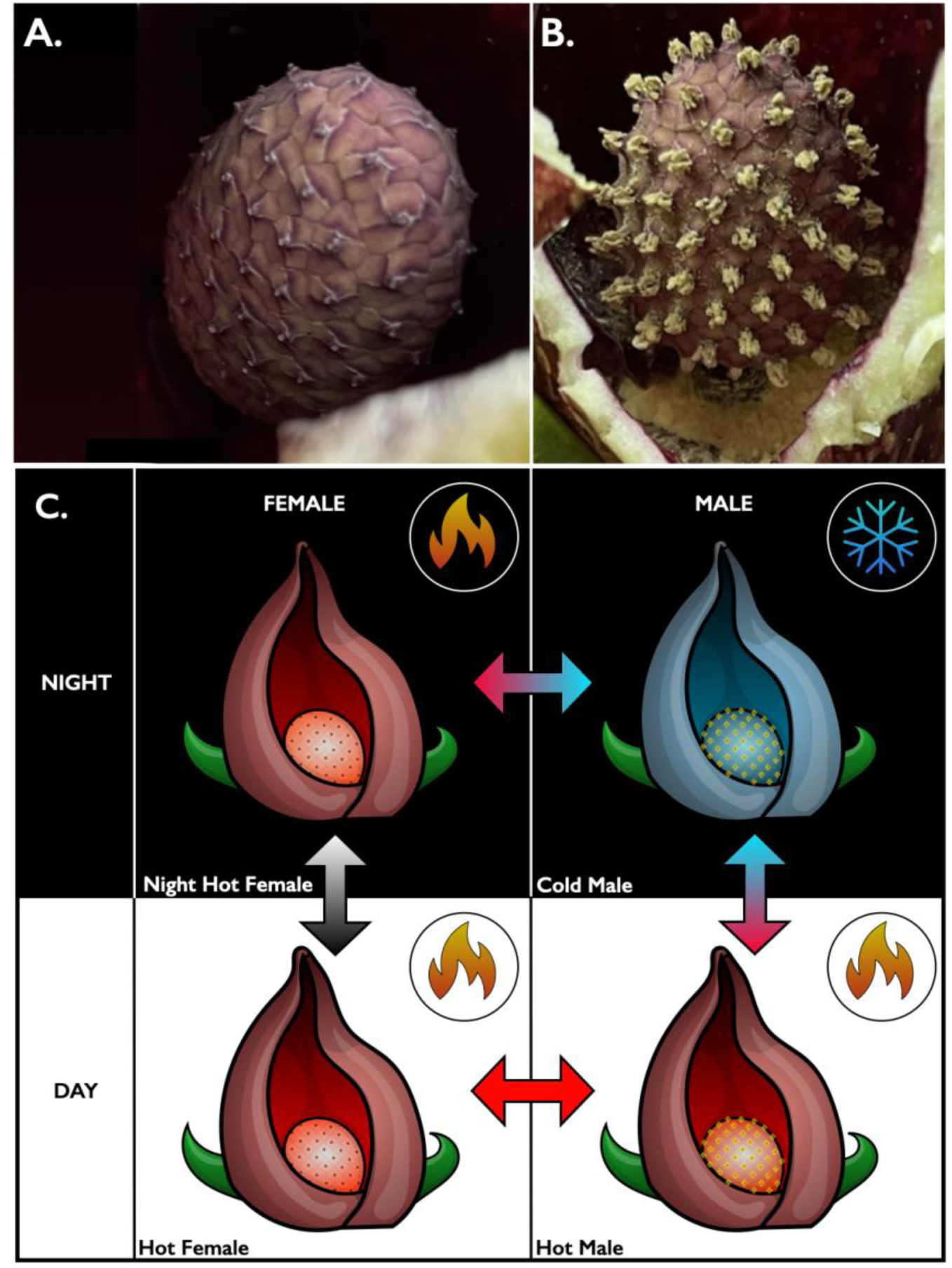
Male (A) and Female (B) spadices of the Eastern Skunk Cabbage. Male and female plants were sampled at midday and midnight.

### Sample preparation

Samples were removed from -80°C and the outer layer of the spadix (i.e., florets) were carefully separated from the pith using a razor blade over liquid nitrogen. The florets were immediately powdered in liquid nitrogen and ∼110 mg was collected for analysis. The samples were grouped according to sex and collection time-of-day, with n=4 in the “Hot Female” group (midday), n=4 in the “Hot Male” group (midday), n=4 in the “Cold Male” group (midnight), and n=4 in the “Night - Hot Female” group (midnight). Notably, we selected samples that best reflected the expected thermal patterns for that sample group. Samples were lyophilized (Labconco Freezone 4.5) and shipped to The Metabolomics Innovation Centre (TMIC; University of Alberta, Canada) on dry ice for processing. The Nova MT Sample Normalization kit was used to determine the total concentrations of the samples. Various volumes of water were added to adjust the samples to 8 mM. Sample normalization was based on total metabolite concentrations measured in all individual samples. Single replicates were measured through 4-channel analysis with 1 labelling kit used per channel.

### Chemical isotope labeling (amine/phenol)

19 μL of LC-MS grade water was added prior to amine-/phenol-labeling of one aliquot. The labeling protocol outlined in the SOP of the kit was then strictly followed. 12.5 μL of buffer reagent (Reagent A) and 37.5 μL of either 12C2-labeling (individual samples and the pooled sample) or 13C2-labeling (pooled sample) reagent (Reagent B) was added to the samples, which were then vortexed, followed by spinning down. The mixtures were incubated at 40 °C for 45 minutes, and then 7.5 μL of quenching reagent (Reagent C) was added to quench the excessive labeling reagent. The mixtures were incubated at 40 °C for another 10 minutes, and then 30 μL of pH adjusting reagent (Reagent D) was added.

### Chemical isotope labeling (carboxyl)

19 μL of LC-MS grade ACN was added prior to carboxyl-labeling labeling of one aliquot. The labeling protocol outlined in the SOP of the kit was then strictly followed. 10 μL of catalyzing reagent (Reagent A) and 25 μL of 12C2-labeling (individual samples and the pooled sample) or 13C2-labeling (pooled sample) reagent (Reagent B) was added to the samples, which were then vortexed, followed by spinning down. The mixtures were incubated at 80 °C for 60 minutes, and then 40 μL of quenching reagent (Reagent C) was added to quench the excessive labeling reagent. The mixtures were then incubated at 80 °C for another 30 minutes.

### Chemical isotope labeling (hydroxyl)

19μL of LC-MS grade ACN was added prior to hydroxyl-labeling of one aliquot. The labeling protocol outlined in the SOP of the kit was then strictly followed. 25 μL of reaction-activating reagent (Reagent A) and 40 μL of 12C_2_-labeling (individual samples and the pooled sample) or 13C2-labeling (pooled sample) reagent (Reagent B) was added to the samples, which were then vortexed followed by spinning down. The mixtures were incubated at 60 °C for 60 minutes, and then 5 μL of quenching reagent (Reagent C) was added to quench the excessive labeling reagent. The mixtures were incubated at 60 °C for another 10 minutes, and then 25 μL of pH adjusting reagent (Reagent D) was added.

### Chemical isotope labeling (carbonyl)

19 μL of LC-MS grade water was added prior to carbonyl-labeling of one aliquot. The labeling protocol outlined in the SOP of the kit was then strictly followed. 25 μL of pH adjusting reagent (Reagent A) and 25 μL 12C_2_-labeling (individual samples and the pooled sample) or 13C_2_-labeling (pooled sample) reagent (Reagent B) was added to the samples, which were then vortexed, followed by spinning down. The mixtures were incubated at 40 °C for 60 minutes, then placed in a -80 °C freezer for 10 minutes to stop the reaction. The solution was then dried under a nitrogen blower, and the metabolites were re-suspended in 100 μL of LC-MS grade ACN/water (50:50 v/v).

### Mixing

12C_2_-labeled individual samples were mixed with 13C_2_-labeled reference samples in equal volume, forming a mixture ready for LC-MS analysis. A quality control (QC) sample was prepared before LC-MS analysis of the entire sample set by mixing equal volumes of a 12C-labeled and a 13C-labeled pooled sample.

### LC-MS analysis conditions

The instrument used for analysis was the Agilent 1290 LC linked to Agilent 6546 QTOF Mass Spectrometer, with the column used being Agilent eclipse plus reversed phase C18 column (150 x 2.1 mm, 1.8 μm particle size). MPA was 0.1% (v/v) formic acid in water, and MPB was 0.1% (v/v) formic acid in acetonitrile with a gradient of t=0min, 25%B; t=10min, 99%B; t=15min, 99%B; t=15.1min, 25% B; t=18 min, 25% B. LC-MS analysis strictly followed the SOP (e.g. Rapid LC-MS Analysis for HP-CIL Metabolomics Platform), and QC samples were injected every 10 sample runs to monitor instrument performance with the following parameters: flow rate = 400 μL/min, column oven temperature = 40 °C, mass range = m/z 220-1000 and acquisition rate = 1Hz.

### Data processing and cleaning

A total of 92 LC-MS data from 4-channel analysis (23 LC-MS data, including 3 QC, in each channel) were exported to a .csv file with Agilent MassHunter software, then exported data were uploaded to IsoMS Pro 1.2.20. Data Processing was performed after Data Quality Check with parameters: minimum m/z = 220, maximum m/z = 1000, saturation intensity = 15000000, retention time tolerance = 9 seconds, and mass tolerance = 10ppm. 4-channel LC-MS data of one sample were combined after data processing. Less commonly detected peak pairs were filtered out to ensure data quality.

23 LC-MS data were assigned to five groups in each channel: 4 data files designated as the “Night-Cold Male” group, 4 data files as the “Night-Hot Female” group, 4 data files as the “Hot Male” group, 4 data files as the “Hot Female” group, and 3 data files as the “QC” group. Peak pairs lacking data in at least 80.0% of samples within any group were filtered out. Subsequently, the data were normalized using the Ratio of Total Useful Signal.

### Metabolite identification

A three-tier ID approach was used to perform metabolite identification. In tier 1, peak pairs were searched against a labeled metabolite library (CIL Library) based on accurate mass and retention time, providing positive identification results with the highest confidence. In tier 2, the remaining peak pairs were searched against a linked identity library (LI Library), providing high-confidence putative identification results based on accurate mass and predicted retention time matches. In tier 3, the remaining peak pairs were searched, based on accurate mass match, against the MyCompoundID (MCID) library for more comprehensive putative results.

The parameters used for metabolite identification: retention time tolerance for CIL Library ID = 10 seconds, retention time tolerance for LI Library ID = 75 seconds, mass tolerance for CIL Library ID = 10ppm, mass tolerance for LI Library ID = 10ppm, and mass tolerance for mass-based database ID = 10 ppm. We also tested for the presences of thyroid hormones (3-Iodo-L-tyrosine; 3,3′,5-Triiodo-L-thyronine; 3,5-Diiodo-L-thyronine) with EIC using QCs in A and C channels.

### Univariate analysis

Pairwise comparisons of identified metabolites were used to determined significant Fold Change (FC) by calculating the average peak ratio value (12C-labeled individual sample vs. 13C-labeled pool) in one study group (Group 1) to that in the other study group (Group 2). Volcano plots were plotted using MetaboAnalyst (Pang et al., 2021), with the processing parameters: Sample Normalization: None; Data scaling: Auto Scaling. Volcano plots were constructed by plotting the fold change (FC) of each metabolite against p-value. The fold change was calculated as Mean(Sample 1) / Mean(Sample 2). In the sex-specific plots, the “Male” samples were treated as Sample 1 and the “Female” samples were treated as Sample 2. In the circadian-specific plots, “Night” samples were treated as Sample 1 and “Day” samples were treated as Sample 2. Plots were specified using FC > 1.2 or < 0.83 and p-value < 0.05 as criteria in accordance with the findings of TMIC Li-Node (2023).

### Filtering significant metabolites

Metabolites identified as having a significant fold change (see above) were cross examined through pairwise comparisons to retain potential thermoregulatory metabolites and ‘filter out’ metabolites with fold changes more correlated with sex and circadian rhythm. Specifically, the “Male” treatment group (Night-Cold Male vs Hot Male) and the “Night” treatment group (Night-Cold Male vs Night-Hot Female) were used to establish metabolite peak pairs exhibiting concentration changes from the heterothermic to the homeothermic state. However, these comparisons also confound circadian rhythm and sex, respectively. Therefore, any metabolites demonstrating similar concentration changes in the “Circadian” analysis (Night-Hot Female vs Hot-Female) were excluded as they were more likely to be regulated by circadian rhythm than thermoregulation. Similarly, any metabolites showing comparable concentration shifts in the “Sex” analysis (Hot Male vs Hot Female) were also filtered out as sex-specific rather than thermoregulatory. An exception was made for peak pairs in the “Circadian” and “Sex” treatments that exceeded a FC > 1.2 or < 0.80 in the same direction as the potential thermoregulatory peak pairs.

### Metabolite Enrichment Analysis

Metabolite enrichment analysis was performed using MetaboAnalyst version 6.0 (Pang et al., 2021) with pathways sourced from RaMP-DB (Relational Database of Metabolic Pathways), which integrates metabolite and lipid pathways from KEGG, HMDB, Reactome, and WikiPathways (Braisted et al., 2023). The RaMP-DB database was selected as it offers a comprehensive collection of 3,694 metabolite and lipid pathways, providing extensive coverage of known biochemical pathways.

The list of filtered metabolites was uploaded to MetaboAnalyst, and the RaMP-DB pathway library was specified for pathway-based enrichment analysis. MetaboAnalyst first mapped metabolites to their respective pathways, leveraging the database’s multi-source integration to enhance annotation accuracy and pathway coverage. Metabolite Set Enrichment Analysis (MSEA) was then applied to identify pathways significantly enriched within the dataset. To correct for multiple comparisons, we applied the Holm-Bonferroni method, producing adjusted p-values to account for potential false discoveries across the numerous pathways analyzed. Enriched pathways were identified based on a significance threshold of adjusted p < 0.05.

### Principal component analysis

Principal Component Analysis (PCA) was performed on filtered metabolites using MetaboAnalyst version 6.0 (Pang et al., 2021). Data were mean-centered and scaled to unit variance prior to analysis to ensure that each variable contributed equally. PCA was applied to reduce the dimensionality of the dataset, allowing for the visualization of sample clustering based on principal components. To assess the significance of group differences, permutational multivariate analysis of variance (PERMANOVA) was applied, as implemented within MetaboAnalyst. PERMANOVA tests were conducted based on Bray-Curtis dissimilarity matrices with 999 permutations, providing a pseudo-F statistic to evaluate whether the observed clustering was greater than would be expected by chance (Anderson, 2001).

## Results

A total of 10,343 unique peak pairs were detected with an average of 10,046 ± 87 peak pairs per sample. Of these, 9230 peak pairs (89.2%) could be positively identified or putatively matched. Among the identified metabolites, 2131 peak pairs (20.6%) were classified as high confidence (tiers 1 and 2). Peak pairs were searched against a labelled metabolite library (CIL Library) based on accurate mass and retention time, resulting in 461 peak pairs positively identified in tier 1. Additionally, 1670 peak pairs were putatively identified in tier 2 using a linked identity library (LI Library) based on accurate mass and predicted retention time matches. The remaining peak pairs were searched against MyCompound ID (MCID) based on accurate mass match. Specifically, 1466 peak pairs matched in the zero-reaction library, 4169 peak pairs matched in the one-reaction library, and 1464 peak pairs matched in the two-reaction library. Notably, we did not detect any thyroid hormones (3-Iodo-L-tyrosine; 3,3′,5-Triiodo-L-thyronine; 3,5-Diiodo-L-thyronine) across our samples.

### Univariate Analysis

#### Metabolites correlated with thermoregulation

Binary comparison of the significant changes in concentrations of metabolites from the Night Cold Male sample group to the Day Warm Male sample group revealed 704 metabolites that demonstrate a significant fold change from midnight to noon. Of these, 256 (36.4%) metabolites displayed a significant increase in concentration and 448 (63.6%) metabolites experienced a significant decrease (Fig. 3a).

**Figure 3.**
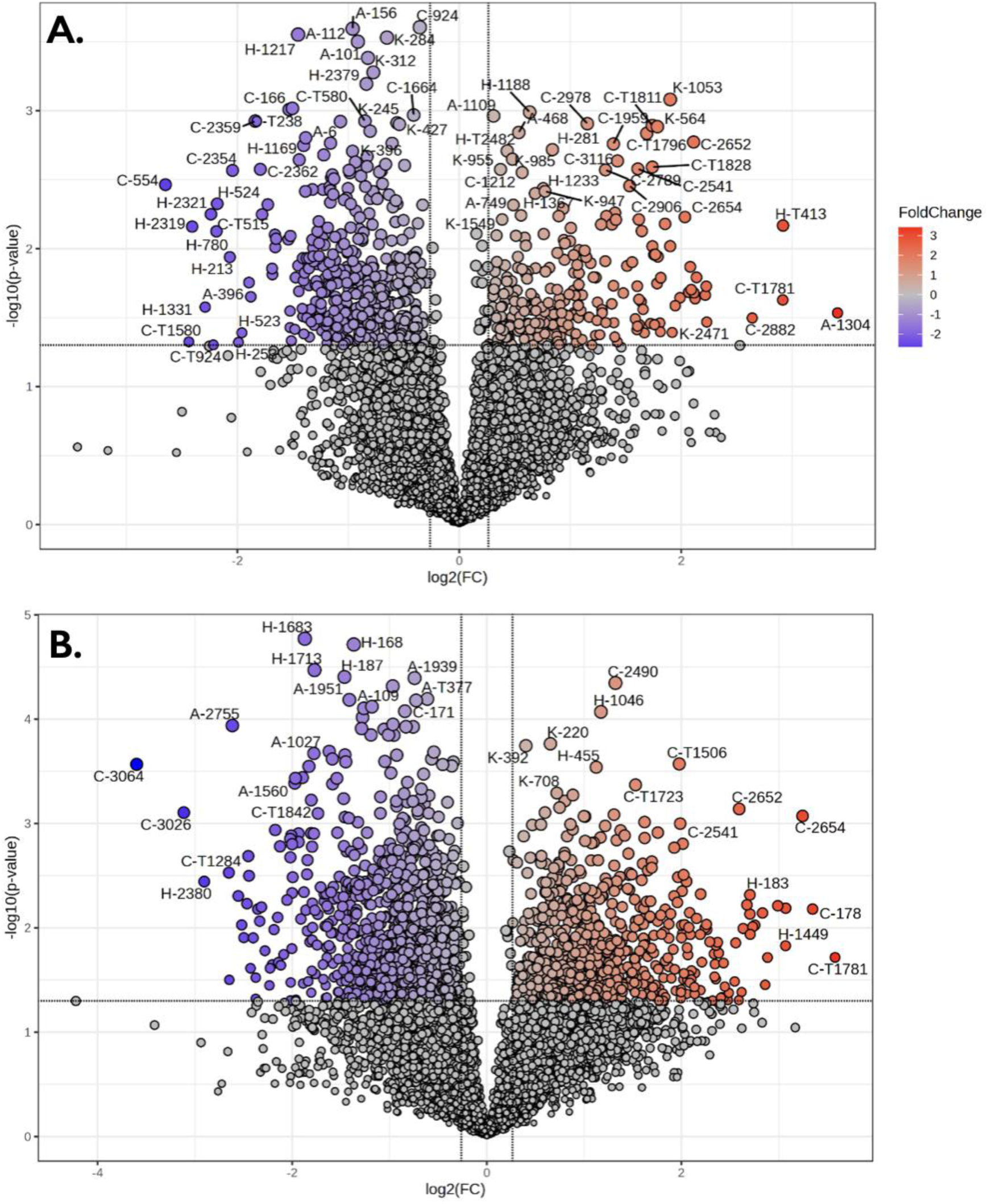
Heterothermic vs homeothermic comparisons showing changes in concentrations of male metabolites sampled from midnight to midday (i.e., Cold Male vs Hot Male; A), and from male to female metabolites sampled at midnight (i.e., Cold Male vs Night-Hot Female; B).

Binary comparison of the significant changes in concentrations of metabolites from the Night Cold Male sample group to the Night Warm Female sample group revealed 1976 metabolites that demonstrate a significant fold change from males at midnight to females at midnight. Of these, 810 (41%) metabolites displayed a significant increase in concentration and 1166 (59%) metabolites experienced a significant decrease (Fig. 3b).

#### Potentially confounding metabolites

Binary comparison of the significant changes in concentrations of metabolites from the Night Warm Female sample group to the Day Warm Female sample group revealed 287 metabolites that demonstrate a significant fold change from midnight to midday. Of these, 104 (36.2%) metabolites displayed a significant increase in concentration and 183 (63.8%) metabolites experienced a significant decrease (Fig. 4a). Notably, these represent metabolites that are likely circadian-regulated since female flowers are homeothermic.

**Figure 4.**
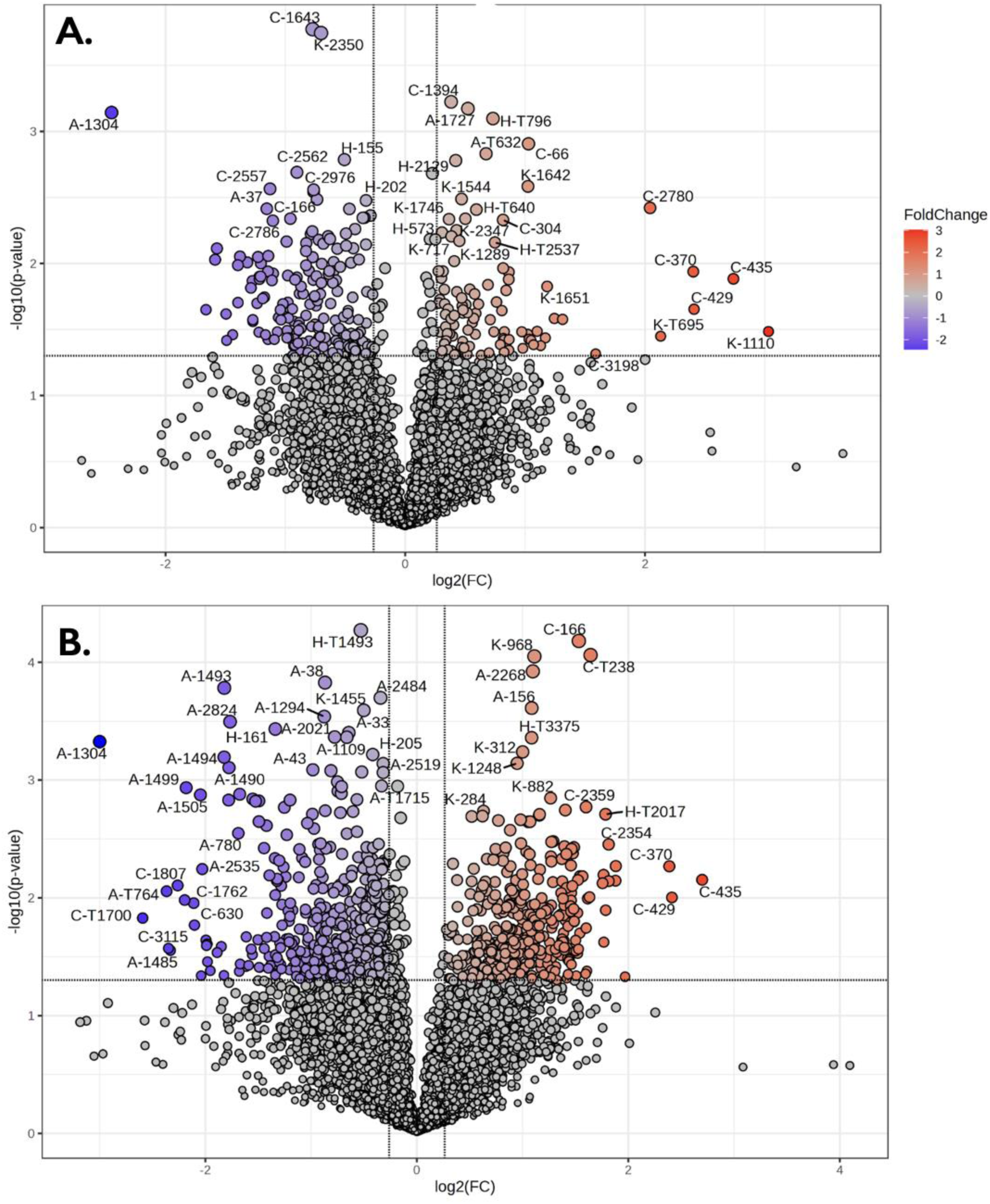
Circadian and sex-dependent comparisons in concentrations of female metabolites sampled from midnight to midday (i.e., Night-Hot Female vs Hot Female; A), and male and female metabolites sampled at midday (i.e., Hot Male vs Hot Female; B).

Binary comparison of the significant changes in concentrations of metabolites from the Day Hot Male sample group to the Day Hot Female sample group revealed 904 metabolites that demonstrate a significant fold change from males at noon to females at noon. Of these, 392 (43.4%) metabolites displayed a significant increase in concentration and 512 (56.6%) metabolites experienced a significant decrease (Fig. 4b). Notably, these represent metabolites that are significantly different between the sexes since both have high thermoregulatory precision at midday.

### Filtering of Significant Metabolite “Hits”

#### Filtering of potential thermoregulatory metabolites

A total of 2680 metabolites changed significantly in Night Cold Male ***vs.*** Day Warm Male, or Night Cold Male ***vs.*** Day Warm Female sample groups. In other words, we identified 2680 metabolites that potentially regulate the circadian phase transition between homeothermy and heterothermy in male spadices or the sexual dichotomy in hetero/homeothermy between males and females at night. Among these metabolites, 254 (9.5%) were shared between the sample groups, indicating potential roles in thermoregulation.

#### Exclusion of circadian rhythm- and sex-specific metabolites

The 287 Metabolites identified to change significantly between Day Warm Female ***vs.*** Night Warm Females were cross-examined with the potential thermoregulatory metabolites in the Day Warm Males ***vs***. Night Cold Males. We excluded metabolite peak pairs that demonstrated similar concentration shifts in both groups (i.e., FC > 1.2 or < 0.80). Of the initial 287 diurnal metabolite peak pairs, 15 (5.2%) were shared, with 1 meeting the exclusion criteria. Similarly, metabolites identified as significant sex-related metabolites (i.e., change between Day Warm Females ***vs***. Day Warm Males) were also cross-examined with potential thermoregulatory metabolites, employing the same exclusion criteria. Among the initial 904 sex-specific metabolite peak pairs, 17 (1.9%) were shared, with 3 metabolites meeting the exclusion criteria. The low overlap between thermoregulatory candidates and circadian- or sex-specific metabolites suggests that the majority of candidate thermoregulatory metabolites are not strongly confounded by these variables.

#### Potential Thermoregulatory Metabolites

Our overlapping list of 254 potentially thermoregulatory metabolites was refined to a pool of 190 (Table S1) by excluding overlapping compounds with different directional changes or those associated with circadian rhythm or sex. However, mass spectrometry profiles of 18 metabolites were unable to be matched to known compounds. Five (2.9%) of these 190 metabolites were identified with a tier 1 level of confidence, 45 (26.2%) were identified with a tier 2 level of confidence, and 122 (70.9%) were identified with a tier 3 level of confidence.

#### Principal Component Analysis

The first principal component (PC1) accounted for 55% of the total variance, while the second principal component (PC2) explained an additional 10.7% (Fig. 5; F-value: 25.534; R-squared: 0.64588; p-value based on 999 permutations: 0.002). Notably, the primary axis of variation predominantly distinguished the cold sample group (i.e., males at midnight; blue ellipsis) from the warm sample groups (i.e., females at midday and midnight, males at midday; pink ellipsis). We therefore used sample loadings for PC1 to determine the top 15% of metabolites (29 total) most positively associated with high thermogenic activity and the 15% of metabolites (29 total) most strongly associated with low thermogenic activity (Table 1; see Table S2 for full PCA loadings list). Of the top 15% of metabolites associated with high thermogenic activity, (R)-4-Phosphopantoic acid, (R)-2,3-Dihydroxy-3-methylpentanoic acid, and maleamic acid were the only compounds identified with high confidence (Tier 2). In contrast, 16 of the 29 compounds most strongly associated with low thermogenic activity were identified with relatively high confidence (Tier 2).

**Figure 5.**
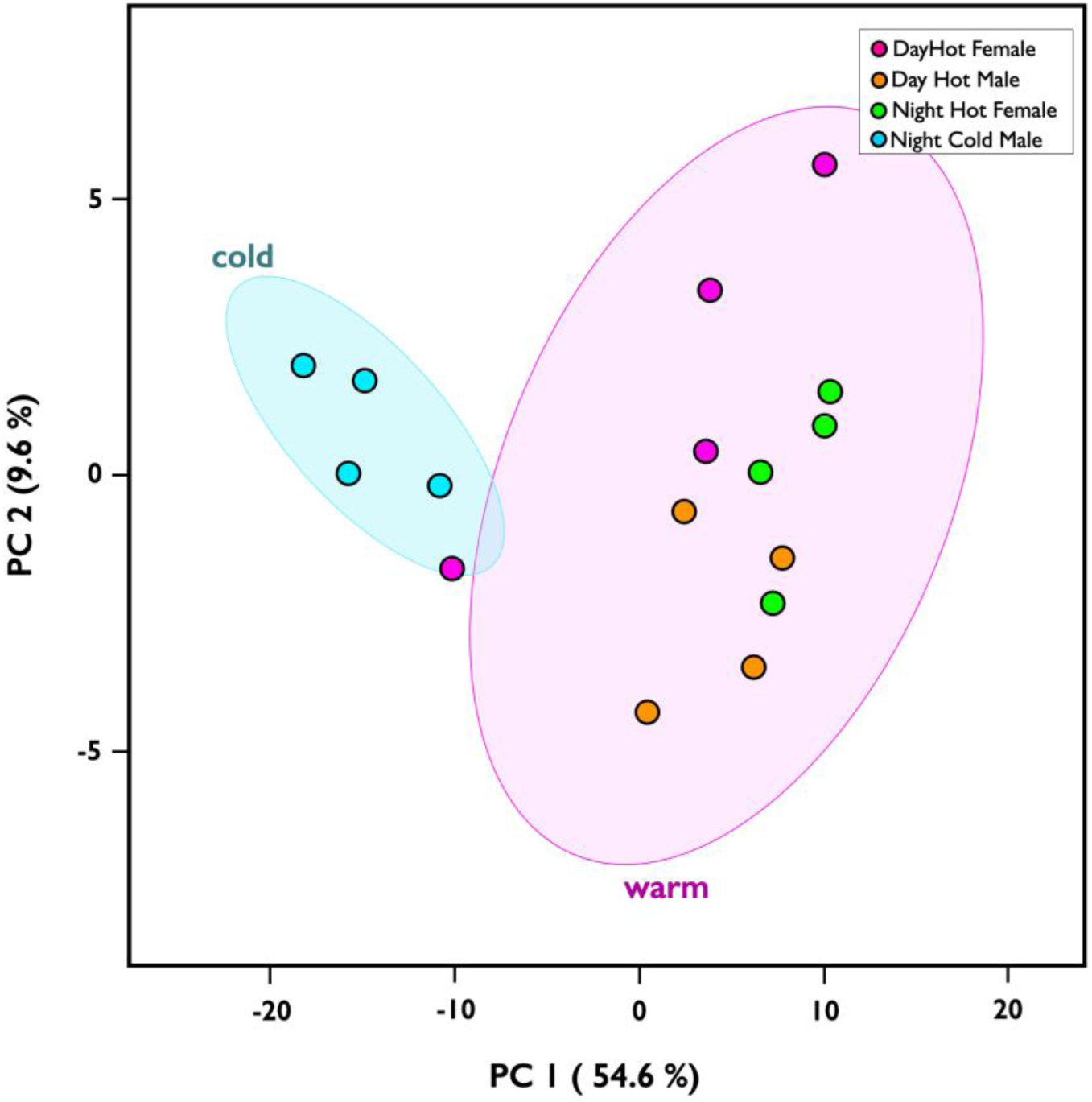
Principal component analysis (PCA) of 190 candidate thermoregulatory metabolites across the four sample groups. PC1 (54.6%) separates thermogenic state, with Night Cold Male spadices (blue; cyan ellipse) clustering distinctly from all warm groups (pink ellipse), which include Day Hot Female (magenta), Day Hot Male (orange), and Night Hot Female (green) samples. PC2 (9.6%) captures residual variance within the warm cluster, partially separating female from male samples. Group differences were significant by PERMANOVA (F = 25.53, R² = 0.646, p = 0.002, 999 permutations).

**Table 1.**
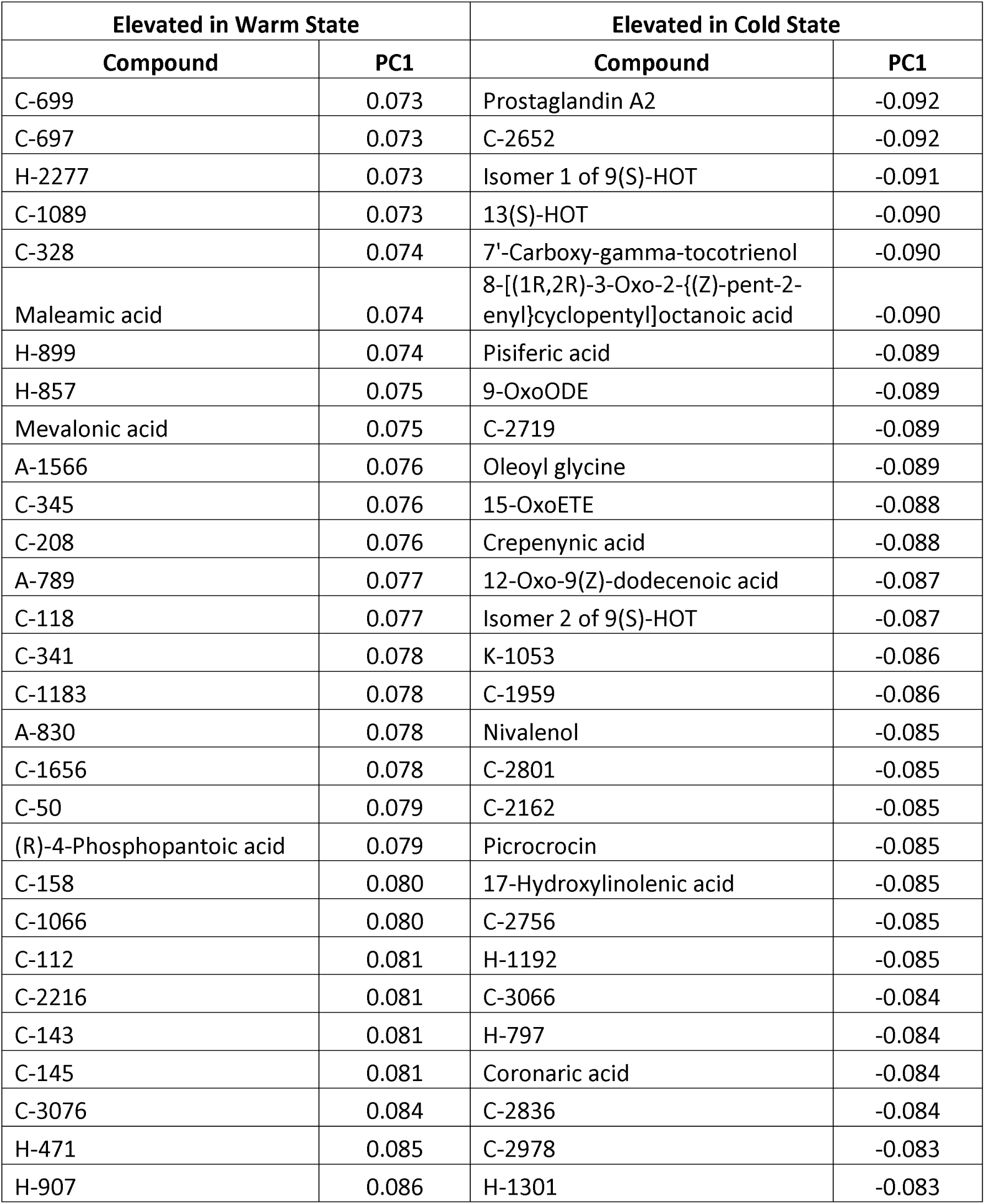
Highest and lowest 15% compound loadings contributing to principal component 1 (PC1).

#### Enrichment Analysis

While only 50 of the 190 metabolites could be identified with relatively high confidence, this list of metabolites was significantly enriched for octadecanoid formation from linoleic acid (Holm P=2.55E-4), linoleic acid oxylipin metabolism (Holm P= 3.17E-4), and eicosanoid metabolism via cyclooxygenases (COX; Holm P= 0.02; Fig. 6).

**Figure 6.**
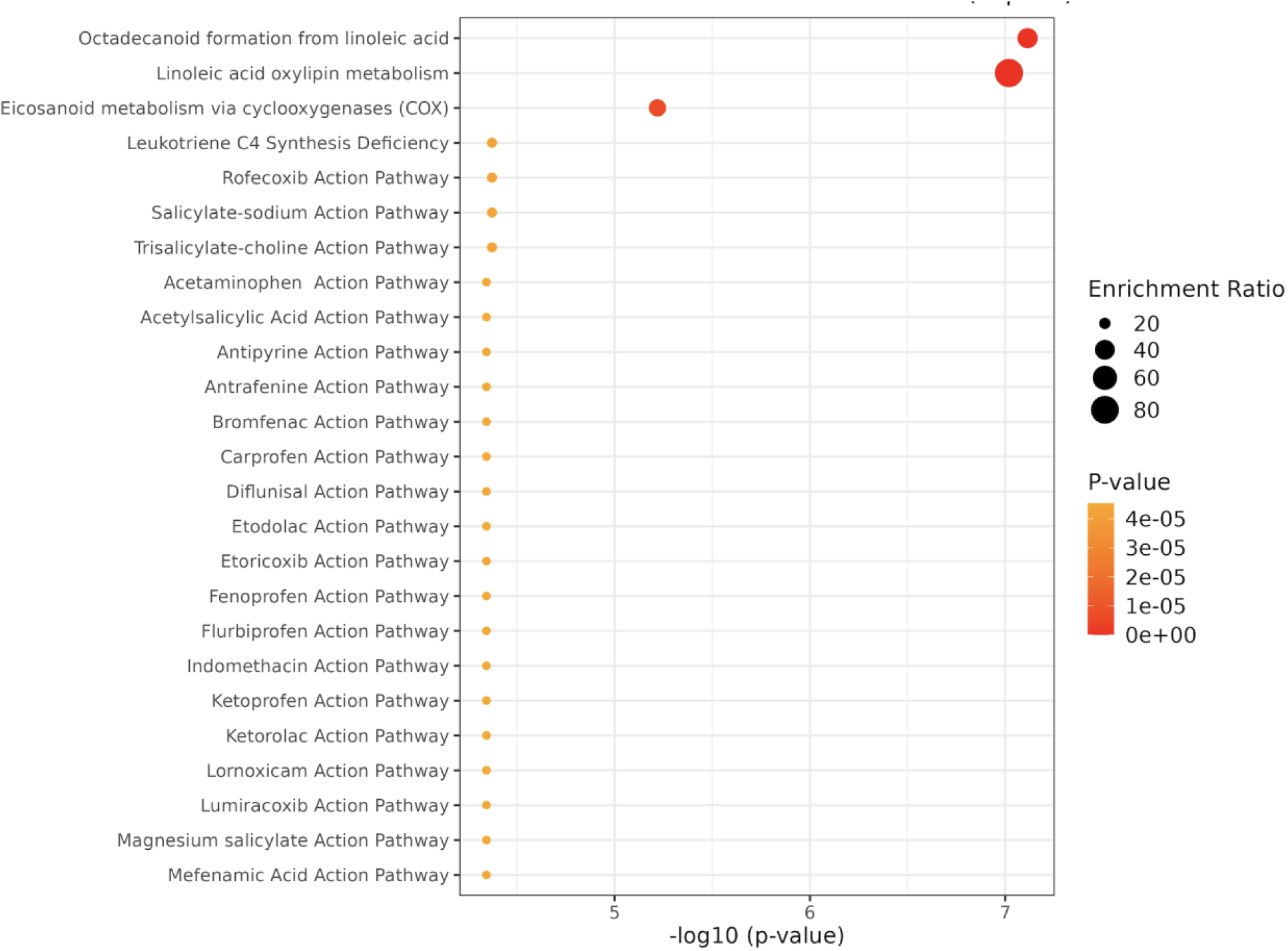
Metabolite Set Enrichment Analysis (MSEA) of 50 high-confidence candidate thermoregulatory metabolites against the RaMP-DB pathway library. The top 25 significantly enriched pathways are shown, ranked by −log10(p-value). Bubble size reflects the enrichment ratio and color indicates p-value (yellow = less significant, red = more significant). The remaining enriched pathways largely represent anti-inflammatory drug action pathways that share oxylipin and eicosanoid targets, consistent with the linoleic acid metabolism theme.

## Discussion

We leveraged the sexual dimorphism in thermogenic strategy of the Eastern skunk cabbage to identify potential signalling pathways that support its remarkable capacity for thermoregulation, the most precise in the plant kingdom. This innovative approach allowed us to refine a set of more than 9000 compounds to a group of 190 potentially thermoregulatory metabolites. Although salicylic acid has been proposed as a regulator of thermogenesis in plants (Raskin et al., 1987; Raskin et al., 1990), our data do not indicate a pattern consistent with thermoregulatory control. Our results do not rule out a role for salicylic acid in remodeling floral tissues for thermogenesis itself, partly by upregulating AOX. Additionally, despite high iodine concentrations in its roots (Marine et al., 1933), we found no evidence that Eastern skunk cabbage florets contain thyroid hormones T_4_, T_3_, or T_2_. Instead, our findings suggest that fatty acid signaling pathways play a central role in thermoregulatory phase transitions in the Eastern skunk cabbage.

### Potential Thermoregulatory Pathways

Many of the identified metabolites are associated with metabolic fuels and signalling pathways in plants and animals. Among these metabolites, we found significant enrichment for pathways involved in octadecanoid formation from linoleic acid, linoleic acid oxylipin metabolism, and eicosanoid metabolism via cyclooxygenases (COXs; Fig 6). While there were no significant differences in mean linoleic acid concentrations between the sample groups (supplemental data), lipooxygenases (LOXs; plants and animals) and cyclooxygenases (COXs; animals) oxygenate fatty acids like linoleic or linolenic acid to generate signalling molecules. This is interesting because linoleic acid has already been shown to inhibit AOX, promote proton leakage via UCPb, and upregulate the cytochrome oxidase pathway in mitochondrial isolates of *S. renifolius* (Onda et al., 2007; Onda et al., 2008; Ito-Inaba et al., 2008). Perhaps ironically, however, linoleic acid was originally tested for its more universal role in UCP induction, as opposed to suspicion of thermoregulatory relevance in this genus. For instance, linoleic acid also uncouples mitochondrial respiration in non-thermogenic plants such as Jerusalem artichokes (*Helianthus tuberosus* L; Paventi et al., 2006) and potatoes (*Solanum tuberosum*; Hourton-Cabassa et al., 2002). Onda et al. (2015) also found genes for ‘metabolism of fatty acids and oxylipin’ among the GO terms associated with thermogenesis in the transcriptome of the arum of Crete (*Arum concinnatum*).

It is intriguing that dietary linoleic acid and its downstream metabolites can also regulate thermogenesis in mammals (Rafael et al., 1988; Leira et al., 2019; Rao et al., 2023; Zhang et al., 2024; Fiehn et al., 2025; Zhang et al., 2026). For instance, cold-exposure enhances oxylipin concentrations in humans (Kulterer et al., 2020) and oxylipins are thought to provide regulatory control of Ucp1-dependent non-shivering thermogenesis in rodents (Maurer et al., 2018; Leiria and Tseng, 2020). A recent study also showed that some linoleic acid-derived oxylipins are potent agonists of transient receptor potential ankyrin1 (TRPA1; Kaneko et al., 2025), a widespread animal thermosensor for noxious temperatures (Saito et al., 2012; Vlachová, 2021). Plants can also synthesize prostaglandins, prostaglandin-like compounds, and jasmonates (Bild, 1978; Panossian, 1987; Groenewald and Van der Westhuizen, 1997; Mueller, 1998; Mueller 2004). Our original dataset identified 8 putative prostaglandins (Tier 2), with 5 prostaglandins (prostaglandin I2, prostaglandin H2, prostaglandin A1 and A2, 6-Ketoprostaglandin E1; Table S1) strongly associated with thermogenic activity in the Eastern skunk cabbage. Intriguingly, several of these prostaglandins have also been linked to thermal responses in mammals (e.g., Takeuchi et al., 1994; Kandasamy and Williams, 1982; Von Bank et al., 2021) and even in a protozoan ancestor (paramecium; Murata et al., 2000). Validation of these hits as true prostaglandins, however, will require a more targeted metabolomic strategy.

### Potential Thermoregulatory Metabolites

A significant challenge in our study was an inability to identify many of the metabolites most positively associated with thermogenic activity (Table 1). Of these compounds, only (R)-4-phosphopantoic acid, (R)-2,3-dihydroxy-3-methylpentanoic acid, and maleamic acid were identified with high confidence. Phosphopantoic acid is a key intermediate in the biosynthesis of coenzyme A (CoA) and pantothenate (vitamin B5) metabolism, which means that it is critical for cellular energy metabolism and fatty acid oxidation. Similarly, high levels of pentadecanoic acid, a lipolysis by-product, align with its role as a metabolic fuel for the TCA cycle but may also represent an antifungal defense (Sujon et al., 2026). Methylpentanoic acid has been linked to floral scent (Ervik and Knudson, 2003), which may help to explain its higher concentrations when spadices are warmed, presumably promoting volatization (Kozen, 2013). Despite their less confident Tier 3 identifications, 4 of the 18 remaining compounds were putatively narrowed down to a single metabolite (pentadecanoic acid, ethanol, guanidinosuccinic acid, and uracil; Table S1).

Itaconic acid, a dicarboxylic acid derived from the TCA cycle, was also among the few identifiable compounds positively associated with thermogenesis (Table S1). While itaconic acid is well studied in animals, only recently was it found to be an endogenous metabolite in plants (Zhang et al., 2025). Emerging evidence reveals that itaconic acid ramps up ATP production through stimulatory effects on the TCA cycle and electron transport system enzymes in litchi (*Litchi chinensis*; e.g., succinate degydrogenase, SDH), simultaneously upregulating antioxidant defenses (Qin et al., 2026). These mitochondrial effects are particularly interesting because they could conceivably upregulate aerobic respiration to support thermogenesis. Itaconic acid may further promote mitochondrial uncoupling by increasing the activity of SDH (not a proton pump) relative to NADH dehydrogenase (pumps 4 H^+^ per electron pair). Intriguingly, itaconic acid has already been linked to enhanced thermogenesis in mammals through its effects on BAT UCP-1, priming it as a potential therapeutic candidate for metabolic disease (Yu et al, 2024; Geng et al., 2026).

Many of the metabolites negatively associated with thermogenic activity are lipid-signaling molecules linked to linoleic acid oxylipins. For instance, 9□oxoODE, isomers 1 and 2 of 9(S)□HOT, 13(S)-HOT, 12□oxo□9(Z)□dodecenoic acid, and eicosatrienoic acid all represent products of linoleic metabolism pathways. Additionally, the tier 3 compounds that were resolved down to a single potential hit were also oxylipins (e.g., 12,13-EpOME, 13(S)-HODE, 9-HODE). Many of these compounds have not been studied in the context of Eastern skunk cabbage or plant thermogenesis. However, several have been linked to thermal responses in other plants and stress responses more broadly. 9-oxoODE concentrations, for instance, increase with heat stress in wheat (Zhao et al., 2024) and soybean (Zhu et al., 2022), though the mechanisms and consequences of its accumulation are not known. 9(S)□HOT, on the other hand, is associated with mitochondrial aggregation and reduction in membrane potential in pathogen-exposed Arabidopsis (*Arabidopsis thaliana*; Vellosillo et al., 2013). Notably, reactive oxygen species (ROS) play important roles in plant signalling and defense during pathogenesis (De Gara et al., 2003; Bolwell and Daudi, 2009; Cameio et al., 2026;) and 9(S)□HOT is thought to enhance ROS signaling by interacting with complex III of the electron transport (Izquierdo et al., 2021). Colder temperatures, such as those of male spadices sampled at midnight, are also known to increase ROS production as flux through the ETS slows (Suzuki and Mittler, 2006; Airaki et al., 2012).

Traumatin (12-Oxo-9(Z)-dodecenoic acid) represents another oxylipin associated with low thermogenic activity in our study, though it is derived from linolenic acid via 13-LOX. There is little research on whether traumatin represents a metabolic regulator in plants, though its beneficial role in wound healing may involve reduction of ROS damage (Jabłońska-Trypuć et al., 2016). Both 17-Hydroxylinolenic acid and 8-[(1R,2R)-3-Oxo-2-{(Z)-pent-2-enyl}cyclopentyl]octanoic acid (a species of OPC-8:0) are precursors in jasmonic acid synthesis (Jimenez-Aleman et al., 2015). While jasmonic acid was identified across our sample groups, its concentrations did not correlate with thermogenic activity (supplemental data). In contrast, however, enrichment for jasmonic acid signalling was associated with thermogenic activity in the arum of Crete (Onda et al., 2015). OPC-8:0 is typically characterized as an inactive precursor of jasmonic acid (Li et al., 2024), but increases in concentration at low temperatures (Li et al., 2025) may actually possess biological activity (Enomoto et al., 2021) and.

The lack of data linking our identified metabolites to thermoregulation in plants is not particularly surprising considering that thermogenic species are rare and do not hold broad economic or agricultural importance. Thermogenesis is, however, particularly well-studied in vertebrates and our findings indicate intriguing points of convergence in thermoregulatory pathways across these two kingdoms of life. For instance, prostaglandin A2 is a specific transactivator of NOR1 (nuclear receptor 4A3; Kagaya et al., 2005), which helps regulate thermogenic capacity of brown adipose tissue in mice (Kanzleiter et al., 2006), potentially via upregulation of UCP1 (Kumar et al., 2008). Many of the other potentially thermoregulatory LOX metabolites identified in Eastern skunk cabbage also represent peroxisome proliferator-activated receptor γ (PPARγ) ligands in mammals (Kumar et al., 2016). 9-oxoODE and 15-OxoETE are known ligands for PPARγ (Egawa et al., 2015; Powell and Rokach, 2015; Ryamn et al., 2017) and promote thermogenesis through UCP1-dependent and -independent mechanisms (Santos et al., 2015; Kaupang and Hansen, 2020; Miranda et al., 2024). 13(S)-HOT and 9(S)-HOT also activate PPARγ (Kumar et al., 2016; Evans et al., 2024), though whether they directly influence thermogenesis has not been explored. N-acyl amino acids can also upregulate mammalian UCP1 expression for thermogenic uncoupling (Gao et al., 2022). Again, however, whether the N-acyl amino acid identified here (i.e., oleoyl glycine) exerts similar effects has not been tested empirically.

### Potential Mechanism for Homeothermy to Heterothermy Transition

One of the most unique aspects of the Eastern skunk cabbage is its sexual dimorphism in thermoregulatory strategy. Our findings suggest that linoleic acid metabolites including oxylipins and similar compounds are elevated particularly during periods of low thermogenic activity (i.e., in cold male florets sampled at midnight). In plants, LOX expression is circadian-regulated in different species and tissues (Nemchenko et al., 2006; Christensen et al., 2013; Zeng et al., 2017; Zhu et al., 2018). It is conceivable that male florets upregulate LOX expression and activity at night, where increased linoleic acid oxylipin metabolism dampens thermogenesis. Intriguingly, lineolate 9S-lipoxygenase (LOX 5) transcript levels in Asian skunk cabbage florets are approximately 4-fold higher in their pre- and post-thermogenic stages than during active thermogenesis (Tanimoto et al., 2024). Similarly, linolenate 13S-lipoxygenase (LOX6) transcript expression in the titan arum (*Amorphophallus titanum*) appendix increases approximately 4-fold over a period of 4 days as the florets transition to their post-thermogenic phases (Zulfiqar et al., 2024). Many of the LOX-derived metabolites associated with low thermogenic activity in the Eastern skunk cabbage (e.g., 9-OxoODE, 15-OxoETE, OCP-8:0) were 3- to 4-fold higher in the cold male spadices sampled at night relative to warm male spadices sampled at midday. Future work should address whether sex-specific circadian regulation of LOX expression triggers the transition from homeothermy in the female phase to heterothermy in the male phase of Eastern skunk cabbage.

### Potential Mechanisms for Thermoregulation

In most plants, metabolic rates are correlated with ambient temperature. However, similar to endothermic animals, florets of the Eastern skunk cabbage upregulate aerobic metabolism as temperatures cool. Birds and mammals finetune thermogenesis by integrating thermal inputs with temperature setpoints in the hypothalamus, whereas plants obviously lack analogous central regulation, making the thermoregulatory precision of the Eastern skunk cabbage particularly remarkable.

A potential mechanism for this rapid and precise thermoregulation may lie in disinhibition. A thermally sensitive inhibitory system would allow thermogenesis to be actively suppressed at optimal temperatures (∼20-24°C) but become progressively less inhibited as temperatures drop, constituting a negative feedback loop where increased thermogenesis warms the system and restores inhibition. The 3- to 4-fold lower concentrations of linoleate- and linolenate-derived oxylipins in homeothermic florets relative to cold male florets could be consistent with such a model, where oxylipin concentrations are titrated by LOX thermal sensitivity to finetune thermogenesis.

Many of the linoleic acid-derived compounds most strongly associated with low thermogenic activity are cyclopentenones (e.g., prostaglandin A2, OPC-8:0, 15-OxoETE), electrophilic five-membered ring enones that reversibly modify dual cysteine residues to inactivate thioredoxin (TRX) activity (Shibata et al., 2003; Gelhaye et al., 2004; Maynard et al., 2021). This is intriguing because TRXo maintains AOX in its reduced and active form in skunk cabbage mitochondria (Umekawa and Ito, 2019), suggesting that elevated cyclopentenone concentrations could suppress AOX-driven thermogenesis via TRXo inhibition, while declining concentrations would relieve this inhibition and activate heat production. While speculative, future work should focus on this notable link between cyclopentenones and AOX-derived heat production.

## Conclusion

Despite over 1.5 billion years of evolutionary divergence, our study suggests convergence in the regulatory outcomes that govern metabolic heat production in thermogenic plants and mammals. However, our findings reveal that metabolites associated with thermogenesis in mammals may have distinct, even opposing, effects in plants, potentially by modulating flux between the thermogenic alternative oxidase (AOX) pathway and the non-thermogenic cytochrome oxidase (COX) pathway. Further resolution of the metabolite identities underlying thermogenic states in Eastern skunk cabbage will yield new insights into the diverse regulatory strategies plants use to control metabolic heat production. These discoveries may ultimately inspire novel approaches to modulating metabolic pathways relevant to human health.

## Acknowledgements

We thank The Metabolomics Innovation Centre acknowledge TMIC and the Grey Sauble Conservation Authority.

